# Endotoxemia-induced inflammation in the absence of obesity decreases anxiety-like and impulsive behavior without affecting learning and memory

**DOI:** 10.1101/2025.05.19.654970

**Authors:** Molly Klug, Eden Crain, Anna Hayes, Mugil Shanmugam, Claire de la Serre, Scott E. Kanoski

## Abstract

Obesity is associated with increased gut permeability, which contributes to a state of chronic low-grade inflammation. Obesity is also linked with altered neurocognitive functions, including impaired learning and memory. Whether these changes are secondary to neuroinflammation vs. other comorbidities associated with obesity is unknown. Here, we modeled the chronic low-grade inflammation that accompanies diet-induced obesity, but in the absence of obesity or consumption of an obesogenic diet. Male rats were implanted with intraperitoneal osmotic minipumps, continuously dispensing either saline (control) or lipopolysaccharide (LPS), an endotoxin produced in the gut that triggers inflammation when in circulation. Immunohistochemistry results revealed that LPS exposure led to neuroinflammation, with an increased number of Ionized Calcium-Binding Molecule 1 (Iba1^+^) cells in the amygdala and hippocampus in LPS rats vs. controls. Given that these brain regions are associated with impulse control, anxiety-like behavior, and learning and memory, we tested whether chronic LPS treatment impacted these behaviors. Interestingly, LPS exposure did not affect hippocampal-dependent memory in the Morris water maze, novel location recognition, or novel object in context memory tests, suggesting that neuroinflammation in the absence of obesity does not induce memory impairments. Further, chronic LPS significantly decreased anxiety-like behavior in the open field test and food impulsivity in an operant-based procedure. LPS animals also had significantly lower corticosterone and melatonin levels when compared to controls, which may contribute to these behavioral outcomes. These results suggest that the low-grade inflammation associated with obesity is not driving obesity-associated memory impairments but does reduce anxiety and food-motivated impulsive responses.

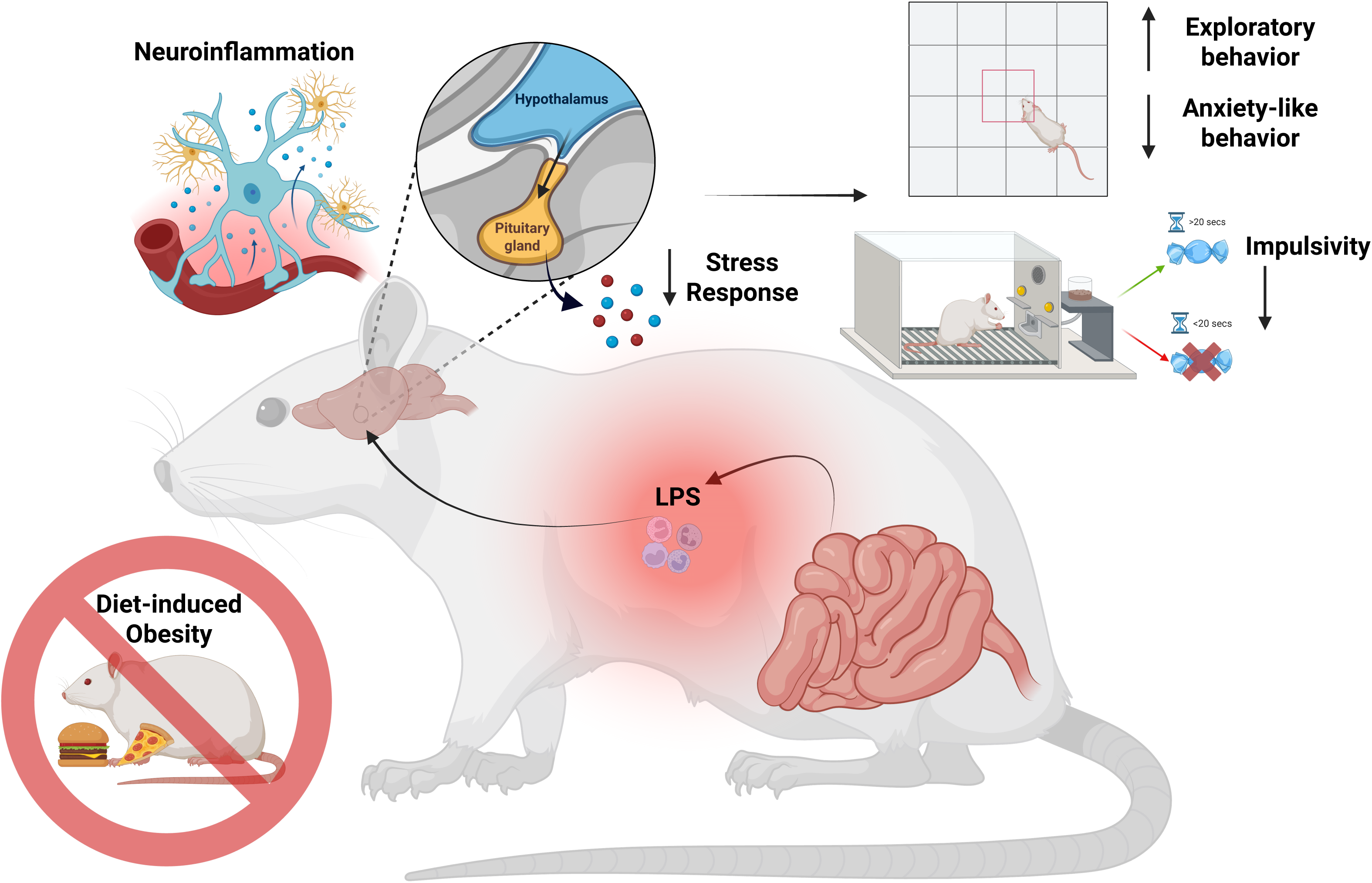

## INTRODUCTION

Obesity is a growing health concern, with approximately 40% of US adults now classified as obese (1). Consumption of a high-fat diet and obesity are each associated with a host of negative health outcomes, including increased gut permeability (i.e., a “leaky gut”) (2–5). One consequence of diet- and/or obesity-associated leaky gut is metabolic endotoxemia, in which higher levels of bacterial-derived lipopolysaccharide (LPS) in circulation contribute to a state of chronic low-grade inflammation (6–8). LPS is a component of the outer membrane of Gram-negative bacteria, which are produced abundantly in the gut and initiate inflammatory responses via Toll-like receptor 4 activation (6, 7). With obesity-associated increases in intestinal permeability (9), LPS leaks into circulation and promotes peripheral inflammation (7, 9). This endotoxemia-induced peripheral inflammation can lead to blood-brain barrier damage and neuroinflammation (10–12), thus representing a putative mechanism through which unhealthy diets and obesity contribute to a range of neurocognitive deficits (13, 14).

Learning and memory processes that require the hippocampus (HPC) are particularly vulnerable to disruption by obesity, and by Western diet (high in fat, sugar, and processed foods) consumption in the absence of obesity (15–20). Both obesity and Western diet consumption are also associated with altered anxiety-like behavior (21), reward-motivated behaviors (22, 23), and impulsivity (24–26), yet these relationships are complex, potentially bidirectional, and poorly understood. Additionally, the neuroinflammation associated with obesity and Western diet consumption can also impact the above cognitive domains (27, 28), further complicating understanding of causal relationships linking obesity with cognitive outcomes.

In addition to neuroinflammation, obesity is linked with various other physiological changes that may impact cognitive function. Disentangling the myriads of possible contributing factors requires isolating them and evaluating cognitive outcomes in the absence of obesity or consumption of a Western diet. Here, we model the LPS endotoxemia-induced chronic systemic inflammation associated with increased gut permeability in otherwise lean rats maintained on a healthy diet. Further, this model induced neuroinflammation in brain regions linked with cognitive processes impacted by obesity, thus offering a platform to evaluate the causal role of endotoxemia-induced neuroinflammation in the absence of obesity on neurocognitive outcomes.

## MATERIALS AND METHODS

### Animals and Housing

Male Wistar rats (∼200g) were obtained from Envigo (Indianapolis, IN, USA) and single-housed in a temperature-controlled 12-hour light/dark cycle facility. Animals were provided ad libitum access to standard rodent chow (Laboratory Rodent Diet 5001, LabDiets, St. Louis, MO, USA) and water. Body weight and food intake were measured at least three times per week. All procedures were approved by the Institutional Animal Care and Use Committees of the University of Georgia and the University of Southern California.

### Surgery

Multiple cohorts were used for this study. For cohort 1, surgeries took place after several days to habituate to the facilities. For cohorts 2 & 3, surgery took place following behavioral training in either the differential reinforcement of low rates of responding (DRL) or progressive ratio (PR) operant tasks. During surgery animals were implanted intraperitoneally with mini osmotic pumps (#2006, Alzet, Cupertino, CA, USA) containing either sterile saline (0.9% NaCl) or an LPS solution providing 12.5 µg/kg/hr (LPS isolated from *Escherichia coli* O26:B6, #L3755, Sigma Aldrich, St. Louis, MO) to be delivered for 6 wks. The LPS dose was based on previous studies (7, 29). To prepare for surgery, rats were shaved on the lower left quadrant of the abdomen and the surgical site was sterilized with betadine. Surgery was performed under isoflurane inhalation anesthesia at a dose of 2-3%. Animals were injected with carprofen or ketoprofen (subcutaneous, 5mg/kg for both) preoperatively for analgesic. A 2cm incision was made and pumps were implanted intraperitoneally, after which the skin and muscle layers were sutured together and closed with sutures or wound clips. Animals were given topical antibiotic post-operatively. Wound clips or stitches were removed 7-10 days post-surgery.

### Behavior testing

Behavior testing started four weeks after surgery as previous research has shown that metabolic endotoxemia is fully developed after four weeks (7). Animals received at least one day of rest between different behavioral tests and behavioral testing was completed by 6 weeks post-surgery. Hippocampal-dependent spatial learning and memory was tested using the Morris water maze (MWM), novel location recognition (NLR), and novel object in context (NOIC) tests. Impulsive responding, food reward effort-based responding, and anxiety-like and exploratory behavior were assessed using the differential reinforcement of low rates of responding (DRL) task (impulsivity), the progressive ratio (PR) reinforcement schedule operant task (food reward motivation), and the Zero Maze (ZM) task and the open field test (OFT) (anxiety-like and exploratory behaviors). A general timeline of experimental procedures is shown in Figure 1A. Behavioral tests were conducted in 3 separate cohorts: Cohort 1 was tested in MWM, NLR, and OFT. Cohort 2 was tested in NOIC, DRL, and ZM. Cohort 3 was tested in PR.

**Figure 1.**
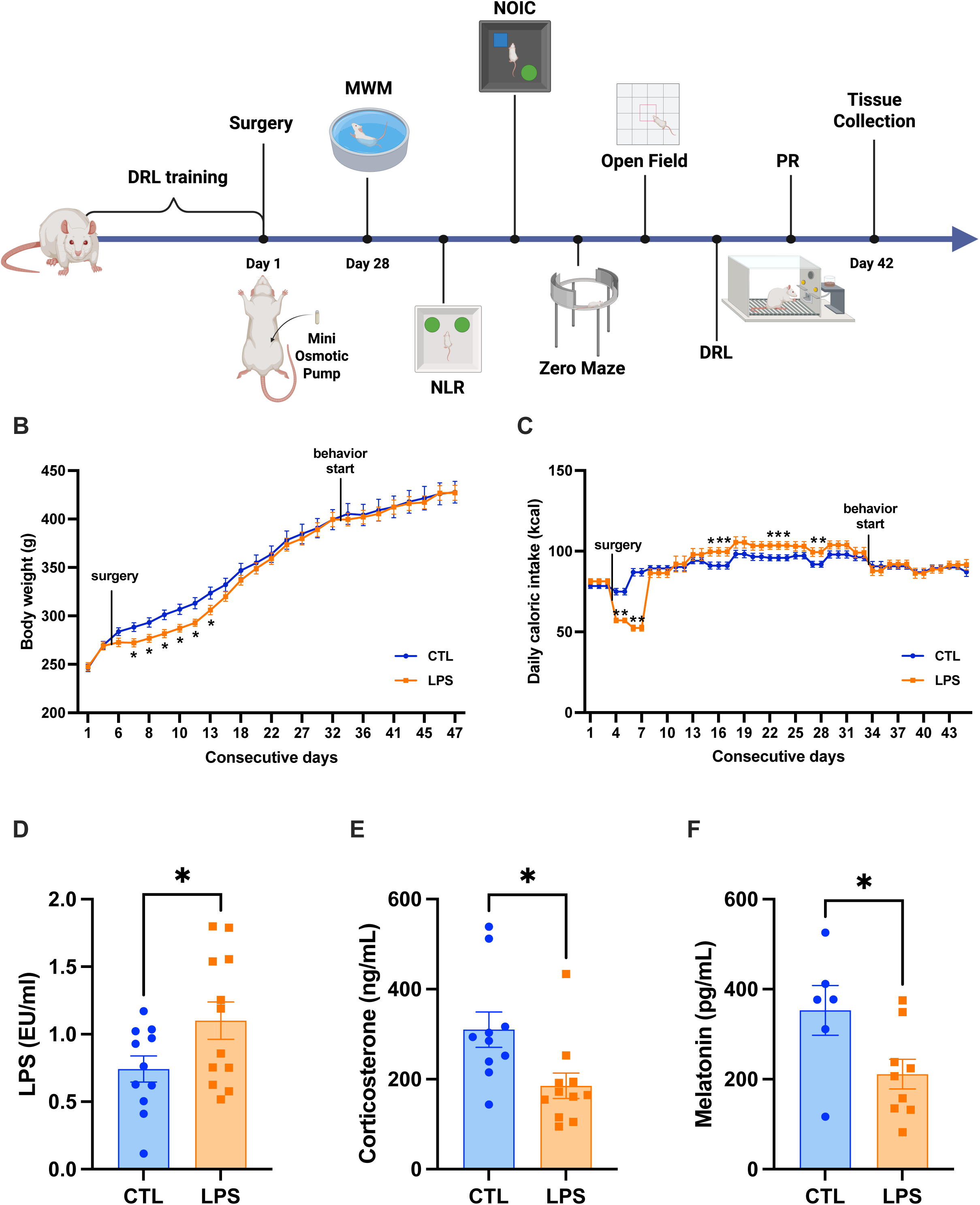
Chronic low-dose LPS administration increases food intake and serum LPS levels, without affecting long-term body weight. **(A)** Experimental timeline (not drawn to scale). Chronic LPS administration causes initial, post-surgical weight loss but does not affect long term body weight **(B).** Similarly, chronic LPS administration causes an initial decrease in food intake, followed by increases in food intake before intake ultimately stabilizes **(C).** Chronic LPS administration increases serum LPS **(D)**, and decreases serum corticosterone (CORT) **(E)** and melatonin **(F)** when compared to controls. (LPS n = 10-12, CTL n = 10-11 for BW, FI, Serum LPS and CORT; LPS = 9, CTL = 6 for melatonin; all between-subjects design; data are means ± SEM; *p<0.05, **p<0.01 ***p< 0.001).

### Morris Water Maze (MWM)

The MWM task, a test of hippocampal-dependent spatial reference memory, was conducted over four days of training with a final memory probe test on day five. Training and testing were done in a blue pool (diameter 170cm) filled with room-temperature water. A resting platform (diameter 10cm) to escape swimming was placed in the center of one quadrant, and large, high-contrast visual cues were placed on the walls surrounding the pool. The platform and cues remained in the same positions for the duration of the experiment. On each training day, animals were placed in the pool and allowed to swim for 60 seconds or until the platform was found. If they had not found the platform in 60 seconds, they were guided to the platform and allowed to stay for 15 seconds. Each rat received five trials per training day with approximately 15-min intervals between trials. After day one, the water was clouded with non-toxic paint and the platform was submerged 1cm below the surface of the water. Rats were started in the same quadrant on days one and two, after which the start quadrant was varied by day. On the test day, the platform was removed and animals were started in the same quadrant as training days one and two. Rats were allowed to swim for 60 seconds before being removed from the pool. The test was recorded with a video camera and analyzed using ANY-maze software to determine swimming patterns and locations (Stoelting Co., Wood Dale, IL).

### Novel Location Recognition (NLR)

The NLR test was conducted using a 50x50x50cm opaque plexiglass arena with patterned markings on the walls to allow the rats to recognize their location. The test was conducted in dim lighting, and light covering was approximately equal between the center and corners. Rats were habituated to the box for 10 min a day for four consecutive days prior to the test day. On test day, two identical objects were placed in parallel corners of the box and animals were allowed to explore for 5 min. After a 5-min interval, animals were returned to the box for 3 min with one of the objects moved to a new location. The object moved was counterbalanced between animals and experimental groups. Boxes and objects were cleaned between animals and trials with 10% ethanol. The test was recorded with a video camera and scored manually by an experimenter blinded to the treatments. For scoring, exploratory behavior was recorded for each object and was defined as follows: nose within 2cm of the object, oriented towards the object or touching the object; not pointing nose into the air, grooming, or sitting on object (unless nose oriented towards object). Exploration was recorded for the initial 5-min trial to assess baseline exploratory behavior and was recorded for the subsequent 3-min trial for each object. Objects were defined as ‘Novel Location Object’ and ‘Familiar Location Object’, and ‘Total Exploration Time’ for the test phase was calculated as the sum of the time spent exploring both the Novel and Familiar Location objects. The ‘Discrimination Index’ was calculated as follows: (time exploring Novel Location Object / Total Exploration Time) x 100%. The test is based on the rodent preference for novelty and assumes that, if the rodents accurately remember the location of the objects, they will favor the novel location object for exploration over the familiar location object. The use of location and contextual cues ensures that this test requires the HPC (30, 31).

### Novel Object in Context (NOIC)

NOIC was conducted to assess HPC-dependent contextual episodic memory (32, 33). The 5-day NOIC procedure was adapted from previous research (17, 18, 34–37). Each day consisted of one 5-min session per animal, with cleaning the apparatus and objects with 10% ethanol between each animal. Days 1 and 2 were habituation to the contexts – rats were placed in Context 1, a semitransparent box (41.9cm L × 41.9cm W × 38.1cm H) with yellow stripes, or Context 2, a black opaque box (38.1cm L × 63.5cm W × 35.6cm H). Each context was presented in a distinct room, both with similar dim ambient lighting yet with distinct extra-box contextual features. Rats were exposed to one context per day in a counterbalanced order per treatment group for the two habituation days.

Following these two habituation days, on the next day each animal was placed in Context 1 containing a single copy of Object A and Object B situated on diagonal equidistant markings with sufficient space for the rat to circle the objects (NOIC day 1). Objects were an assortment of hard plastic containers, tin canisters (with covers), and the Original Magic 8-Ball (objects were distinct from what animals were exposed to in NLR). The sides where the objects were situated were counterbalanced per rat by treatment group. On the following day (NOIC day 2), rats were placed in Context 2 with duplicate copies of Object A. The test day (NOIC day 3) took place one day after NOIC day 2, during which rats were again placed in Context 2, but with single copies of Object A and Object B; Object B was not a novel object per se, but its placement in Context 2 was novel to animals. Rats were consistently placed with their head facing away from both objects when placed into the training and test contexts. On NOIC days 1 and 3, object exploration, defined as the rat sniffing or touching the object with the nose or forepaws, was quantified by hand-scoring of videos by an experimenter blinded to the animal group assignments. The discrimination index for Object B was calculated for NOIC days 1 and 3 as follows: time spent exploring Object B (the “novel object in context” in Context 2) / [time spent exploring Object A + time spent exploring Object B]. Data were then expressed as a percent shift from baseline as: [day 3 discrimination index – day 1 discrimination index] × 100. Rats with intact HPC function will preferentially explore the “novel object in context” on NOIC day 3, while HPC impairment will impede such preferential exploration (32, 33).

### Differential Reinforcement of Low Rates of Responding task (DRL)

As previously described (38), non-food restricted rats were trained in the early nocturnal phase on an operant lever-pressing task to obtain a single, high-fat high-sugar 45mg pellet (HFHS; F05989, Bio-serv; 35% kcal fat and sucrose-enriched) for each press. Training and test sessions were conducted in operant conditioning boxes (Med Associates Inc, St. Albans, VT). After progressively increased delay periods (0-, 5-, 10-second delays, 5 days of training for each) over four weeks of training, rats are given a ‘DRL-2o’ schedule for ten days in which they must wait 20 seconds between lever presses. A lever press prior to conclusion of the 20 second interval resets the clock, is not reinforced, and is considered an impulsive response. Training took place prior to the implantation of the mini osmotic pump, to ensure that learning of the DRL task was unaffected by experimental treatment. After surgery, rats were trained 2x/week, separated by at least one day, at the DRL20 schedule. A single test day took place at the DRL20 schedule five weeks after the pump implantation.

### Progressive Ratio task (PR)

Rats were trained to lever press for 45mg pellet (HFHS; F05989, Bio-serv; 35% kcal fat and sucrose-enriched) in operant conditioning boxes over the course of 6 days, with a 1hr session each day. The first 2 days consisted of fixed ratio 1 with <u>autoshaping</u>, wherein animals would receive 1 pellet for each correct lever press (FR1), and a pellet would dispense automatically every 10 min. The next 4 days consisted of FR1 without autoshaping (2 days) and FR3 (2 days), wherein animals would receive 1 pellet for every 3 presses on the active lever. Training took place prior to the implantation of the mini osmotic pump, to ensure that learning of the task was unaffected by experimental treatment. Post surgery, animals were periodically trained at a FR3 reinforcement schedule. Testing took place roughly five weeks after pump implantation. On the test day rats were placed back in the operant chambers to lever press for HFHS pellets under a progressive ratio reinforcement schedule. The response requirement increased progressively using the following formula: F(i) = 5ê0.2i-5, where F(i) is the number of lever presses required for the next pellet at i = pellet number and the breakpoint was defined as the final completed lever press requirement that preceded a 20-min period without earning a reinforcer, as described previously (39, 40).

### Zero Maze (ZM)

Following established procedures (18), the zero maze apparatus was used to examine exploratory and anxiety-like behavior. The apparatus consisted of an elevated circular track (11.4cm wide track, 73.7cm height from track to ground, 92.7cm exterior diameter) that is divided into four equal length segments: two sections with 3-cm high curbs (open) and two sections with 17.5-cm height walls (closed). Ambient lighting was used during testing. Rats were placed in the maze on an open section of the track and allowed to roam freely for 5 min. The apparatus was cleaned with 10% ethanol between rats. The total distance travelled and time spent in the open segments of the apparatus were measured via video recording using ANY-maze activity tracking software.

### Open Field Test (OFT)

The OFT to assess exploratory and anxiety-like behavior was conducted using a 60x60x40cm clear plexiglass box in brightly lit (∼200 lux) conditions. The arena was divided into nine 20x20cm sections. Animals were placed in the box for 10 min, and the box was cleaned between animals with 10% ethanol. The test was recorded with a video camera and video footage was analyzed for the first 2 min using ANY-maze software. For analysis, the center zone was defined as the 20cm square in the middle of the arena, and corner zones were defined as each of the four corners. Behavior was analyzed for total distance, time in the center zone and corner zones, and number of center zone entries.

### Serum Corticosterone, LPS, and Melatonin

Serum LPS and corticosterone levels were measured in Cohort 1, and serum melatonin was measured in Cohort 2. For serum corticosterone (CORT) and LPS measurements, animals were anesthetized at six weeks post-surgery via CO2 inhalation. Blood was collected via cardiac puncture, allowed to clot on ice for 30 min, and centrifuged for 10 min at 4°C. The supernatant was then flash frozen on dry ice and stored at –80°C until analysis. CORT levels were measured using a Corticosterone ELISA kit (ab108821, Abcam, Cambridge, UK) according to the manufacturer’s instructions.

Serum LPS was measured using the Limulus Amoebocyte Lysate (LAL) method according to established protocols (29). For the LPS measure, all procedures were conducted using endotoxin-free supplies, and surfaces were thoroughly cleaned with 70% ethanol. Briefly, samples were thawed to room temperature, vortexed for one minute, diluted 1:5 with pyrogen free water (LAL Reagent Water, Lonza, Basel, Switzerland), and vortexed for another minute. Samples were held in a 70°C water bath for 10 min, cooled, and diluted 1:5 again for a final dilution of 1:10. Standards were prepared using Control Standard Endotoxin (#EC010-5, Associates of Cape Cod, Inc, East Falmouth, MA, USA) and pyrogen free water. Lastly, a Pyrochrome solution was made by adding Glucashield Buffer to Pyrochrome Lysate Mix (CG1500-5, Associates of Cape Cod, Inc). Samples and standards were plated in an endotoxin-free 96-well plate (Pyroplate, Associates of Cape Cod, Inc). The Pyrochrome solution was added to each well, and the plate was covered in foil and placed on a heat block at 37°C for 75 min. The absorbance was read at 405nm every 5 min to obtain the optimal standard curve with a SpectraMax M3 spectrophotometer (Molecular Devices, San Jose, CA, USA). At 75 min, 50% acetic acid was added as the stop solution and absorbances were read one final time.

For serum melatonin analyses, animals were anesthetized with an intramuscular injection of a cocktail of ketamine (90.1 mg/kg body weight), xylazine (2.8 mg/kg body weight), and acepromazine (0.72 mg/kg body weight), and then rapidly decapitated.

Trunk blood was collected and allowed to clot on ice for 30 min, before being centrifuged for 10 min at 4°C. The supernatant was then flash frozen on dry ice and stored at –80°C until analysis. Melatonin levels were determined using the MILLIPLEX® MAP Rat Stress Hormone Magnetic Bead Panel (RSHMAG-69K; MilliporeSigma, Burlington, MA, USA) by following the manufacturer’s protocol. The standards and samples, diluted at 1:4 with assay buffer, were freshly prepared and run in duplicate. The Luminex plate was read on Bio-Rad Bio-Plex 200 Systems with Manager Software (Hercules, CA, USA) for data acquisition and analysis.

### Immunohistochemistry (IHC) and Neuroinflammation Quantification

IHC analyses were conducted in Cohorts 1 and 2. Animals were anesthetized at six weeks post-surgery via CO2 inhalation and were transcardially perfused with PBS and 4% PFA in PBS. Whole brains were collected and post-fixed in 4% PFA in PBS on ice for 2 hours. Brains were transferred to 30% sucrose, 0.1% NaN3 in PBS for at least 24 hours. Brains were then frozen at –80°C until further analysis. Brains were coronally sectioned at 20µm thickness using a Leica CM1900 cryostat and mounted on negatively charged slides. Sections were collected between bregma 3.2 and 2.2 mm for prefrontal cortex and - 2.3 to - 5.6 for amygdala and hippocampus. Tissue sections were stained for ionized calcium binding adaptor molecule-1 (Iba1), a marker of activated microglia and macrophages, and glial fibrillary acidic protein (GFAP), which is expressed by astrocytes and upregulated in reactive astrocytes. Sections were permeabilized with sodium borohydride and incubated overnight with primary antibody against Iba1 (anti-IBA-1 rabbit, Wako, Cat#019-19741) and GFAP (anti-GFAP rabbit, Abcam, AB7260). Subsequently sections were incubated for 2 hours with Alexa 488 secondary antibody (Invitrogen, #A11008). Slides were examined using a Keyence BZ-X800 fluorescent microscope (Keyence, Osaka, Japan) at 20x magnification. Images were analyzed using QuPath software (41) for cell counts.

IHC to obtain representative images for Figure 2 were conducted using slightly different protocols than quantification, as the two procedures were conducted in different laboratories. For representative images, brains were sectioned at 30µm thickness on a freezing microtome. Sections were collected in 5 series and stored in antifreeze solution at −20°C until further processing. General fluorescence IHC labeling procedures were performed. The following antibodies and dilutions were used: rabbit anti-Iba1 (1:1000, Wako, Cat. # 019-19741) and chicken anti-GFAP (1:2000, Cat. # ab4674, Abcam, Cambridge, UK). Antibodies were prepared in 0.02 M potassium phosphate-buffered saline (KPBS) solution containing 0.2% bovine serum albumin and 0.3% Triton X-100 at 4°C overnight. After thorough washing with 0.02 M KPBS, sections were incubated in secondary antibody solution. All secondary antibodies were obtained from Jackson ImmunoResearch and used at 1:500 dilution at 4°C, with overnight incubations (Jackson ImmunoResearch; West Grove, PA, USA). Sections were mounted and cover slipped using 50% glycerol in 0.02 M KPBS and the edges were sealed with clear nail polish. Photomicrographs were acquired using either a Nikon 80i (Nikon DS-QI1,1280X1024 resolution, 1.45 megapixel) under epifluorescence or darkfield illumination. Images were analyzed using ImageJ (42) for cell counts.

**Figure 2.**
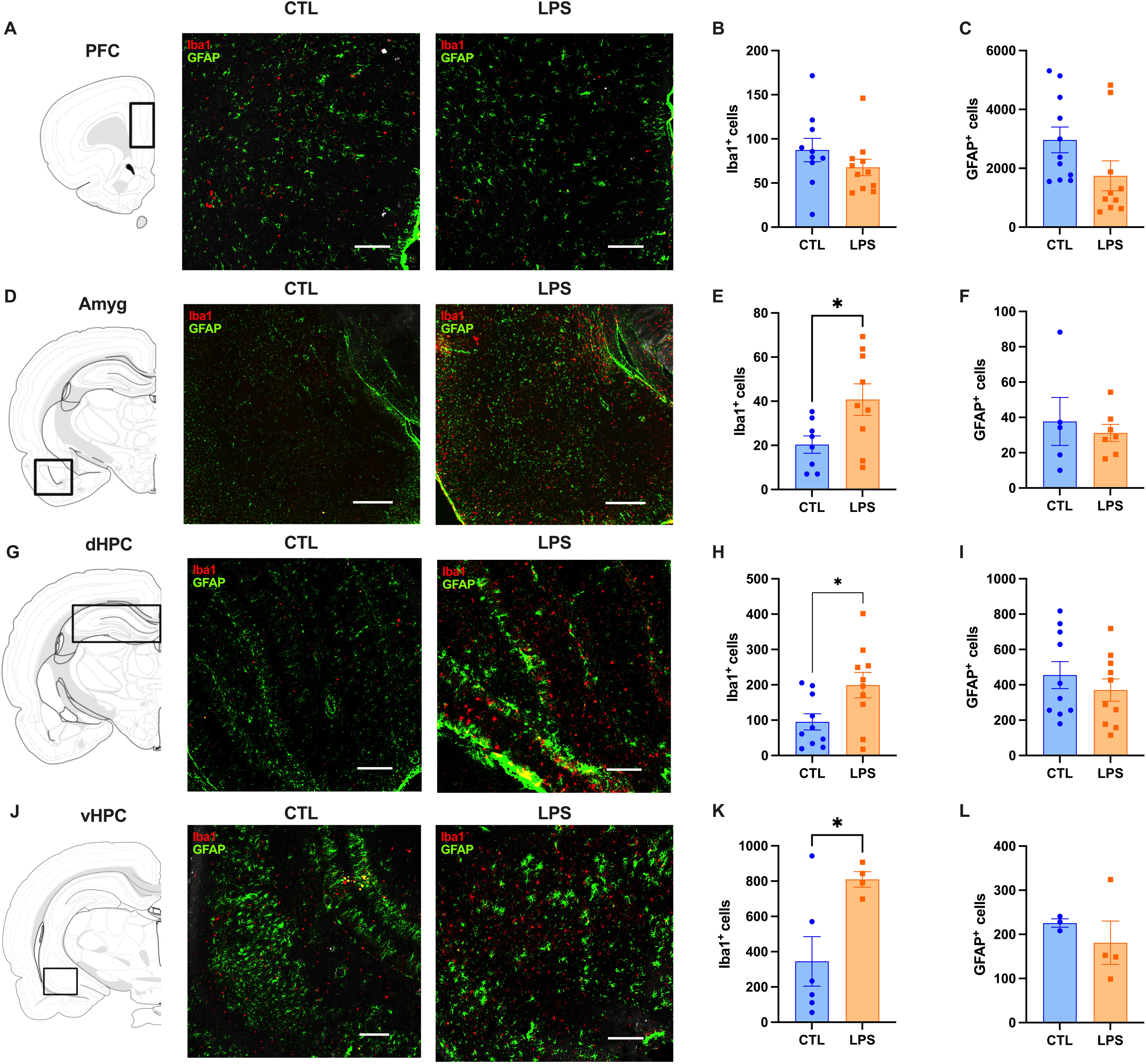
Chronic LPS administration induces neuroinflammation in brain regions associated with learning and memory, impulse control, and anxiety-like behavior. **(A)** Representative images for the prefrontal cortex (PFC), showing control (left) and LPS (right) (Scale bar, 200um). Chronis LPS administration does not affect Iba1 or GFAP staining in the PFC **(B,C)**, nor does it affect GFAP staining in the amygdala (Amyg), dorsal or ventral hippocampus (dHPC; vHPC) **(F, I, L)**. However, Iba1 staining was significantly increased in the Amyg, dHPC, and vHPC **(E, H, K)** in animals receiving chronic low-dose LPS administration. Representative images for the amygdala, dorsal and ventral hippocampus showing control (left) and LPS (right) are shown as well **(D, G, J)** (Scale bar, 200um). (LPS n = 4-10 CTL n = 3-11 ; all between subjects design; data are means ± SEM; *p<0.05).

### Statistical analysis

All statistical analyses were conducted using GraphPad Prism 10.0.3 (GraphPad Software, Inc. Boston, Massachusetts, USA). Data are reported as mean ± SEM. T-tests were used to analyze MWM probe test, NLR, NOIC, OFT, DRL, PR, serum LPS, CORT and melatonin, and immunohistochemistry results. Mixed effects analysis followed by Fisher’s LSD post-hoc analysis for multiple comparisons was used to analyze body weight, food intake, and MWM training. Statistical significance was set at a p-value ≤ 0.05.

## RESULTS

### Chronic LPS administration reduces serum corticosterone and melatonin without influencing long-term food intake or body weight

There were no differences in body weight between groups at the start of the experiment. Analysis of body weight showed an expected main effect of time (F (1.425, 29.50) = 662.8, p < 0.0001) and a significant interaction between time and treatment (F (24, 497) = 2.547, p < 0.0001) (Fig 1B). In both groups, body weight increased with time (Fig 1B). Following surgery and pump implantation, the LPS group lost significantly more weight than the control group (days 7-13, p < 0.05), which corresponds to the beginning of LPS release (Fig 1B). However, following recovery (one-week post-surgery), there were no significant differences in body weight between groups (Fig 1B). There was an overall effect of time (F (5.314, 110.6) = 46.50, p < 0.0001) on food intake, with both groups decreasing intake following surgery. We also observed a significant time by treatment interaction (F (11, 229) = 18.84, p < 0.0001) (Fig 1C). LPS-treated rats exhibited significantly lower food intake compared to control rats following surgery (days 4-7) which corresponds to the beginning of LPS release (p < 0.0001), followed by a period of significantly increased intake (p<0.05) (Fig 1C). However, at the time of behavioral testing, LPS-treated animals did not significantly differ from their saline counterparts in either body weight or food intake (Fig 1C).

As expected, serum LPS was significantly higher in LPS-treated animals compared to control animals (p < 0.05) (Fig 1D). Results also reveal that LPS-treated animals had significantly higher levels of serum corticosterone (p< 0.05; Fig 1E) and melatonin (p < 0.05; Fig 1F) compared to their saline counterparts, suggesting a potentially altered stress response in these animals.

### Chronic LPS administration induces neuroinflammation in brain regions associated with anxiety, impulsivity, and learning and memory

Microglial recruitment and astrocytic reactivity were measured using Iba1 and GFAP immunostaining, respectively, with representative images of regions of interest shown in Figure 2 (A, D, G, J). We analyzed expression in the dorsal hippocampus (dHPC), ventral hippocampus (vHPC), prefrontal cortex (PFC), and amygdala (Amyg), as these are brain regions involved in learning and memory (18), impulsivity (38), and anxiety like behavior (43). Notably, obesity is linked with impaired learning and memory, as well as altered anxiety and impulsivity (15, 21, 26). Results revealed that there were significantly more Iba1^+^ cells in the Amyg (p < 0.05; Fig 2E), dHPC (p < 0.05; Fig 1H), and vHPC (p < 0.05, Fig 2K) of LPS-treated animals compared to controls, while there were no differences between groups in the PFC (p = 0.6, Fig 2B). There were no observed differences in GFAP staining in any of the regions of interest that we examined (Fig 2C, F, I, L). Taken together, these results suggest that chronic low-grade inflammation drives neuroinflammation by increasing microglial activity in brain regions associated with neurocognitive processes.

### Chronic LPS administration has no effect on hippocampal-dependent learning and memory

Due to the increased neuroinflammation in the dHPC and vHPC of LPS-treated animals, and the strong connection between hippocampal-dependent memory impairments and obesity (15, 16, 18), we tested animals in a variety of hippocampal-dependent memory tasks. During the Morris water maze (MWM) task, the amount of time to find the platform (escape latency) was measured for each of 5 trials on 4 consecutive training days to assess hippocampal-dependent learning. The average daily escape latency was calculated for each animal. There was an overall effect of time (F (2.686, 54.62) = 32.39, p < 0.0001), with average escape latency decreasing over time, indicating that rats in both groups learned to find the platform more quickly as training progressed (Fig 3B). However, there were no differences in performance between groups (Fig 3B). Memory was also assessed during a probe test, and no differences between groups in time spent in the platform quadrant (Fig 3C) or total distance travelled were observed (Fig 3D).

**Figure 3.**
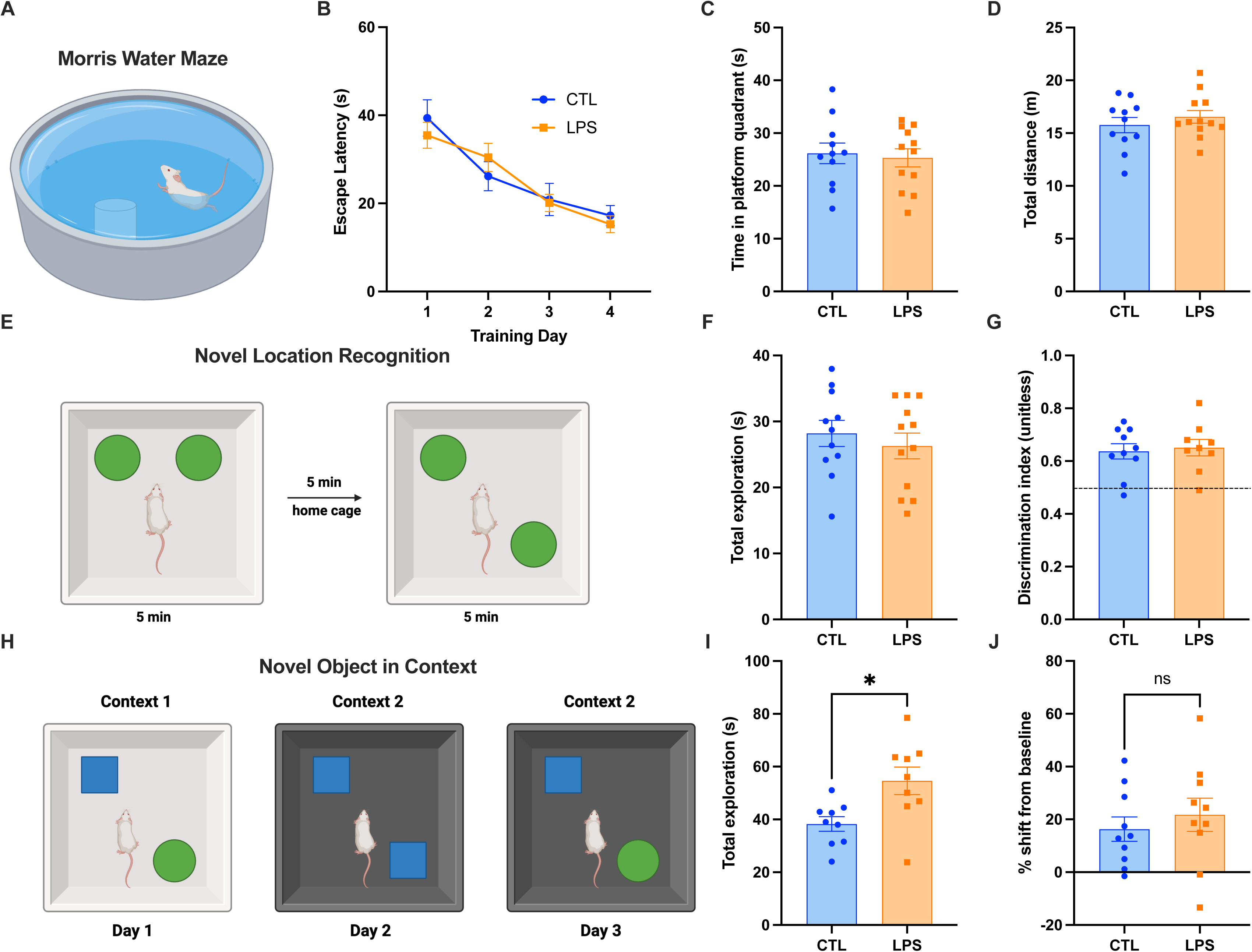
Chronic LPS administration does not affect hippocampal-dependent memory function. Hippocampal-dependent learning and memory was unaffected by chronic low-dose LPS administration in either the Morris water maze (MWM) **(A-D),** the novel location recognition (NLR) **(E-G)**, or the novel object in context (NOIC) tasks **(J).** Total exploration of both objects was significantly increased with chronic LPS administration in the NOIC task **(I)** (LPS n = 12, CTL n = 11 for MWM, NLR; LPS n = 10, CTL n= 10 for NOIC; between-subjects for all experiments; data are means ± SEM; *p<0.05).

We also used the novel location recognition (NLR) and novel object in context (NOIC) tasks to assess hippocampal-dependent learning and memory. For NLR, there were no differences between groups in exploratory behavior (Fig 3F) or discrimination for the novel object (Fig 3G). However, in the NOIC task, LPS-treated animals spent significantly more time exploring both objects when compared to controls (p < 0.05; Fig 3I), though there were no differences in the preference for the novel object as measured by percent shift from baseline (Fig 3J). Together, these data suggest that obesity- and Western diet-associated memory impairments from previous reports are unlikely to be driven by chronic low-grade inflammation or neuroinflammation, or at least not that induced by LPS-mediated endotoxemia.

### Chronic LPS administration reduces anxiety-like behavior and impulsive responding for palatable foods, independently of food motivation

While learning and memory are unaffected by chronic LPS administration, the brain regions with increased microglial activity shown in Figure 2 are also associated with impulsive responding (38) and anxiety-like behavior (43). Given these previous findings and the significant difference in exploratory behavior between LPS-treated and control animals in the NOIC task, animals were tested in the open field test (OFT) and zero maze (ZM) test to assess exploratory and anxiety-like behavior.

Behavior during the first 2 min of the OFT (Fig 4A) was measured to capture the initial response to a new environment. There were no differences between groups in total distance travelled (Fig 4B). However, LPS-treated rats had significantly more entries into the center zone and spent significantly more time there when compared to control animals (p < 0.05; Fig 4C, D), indicating a decrease in anxiety-like behavior and an increase in exploratory behavior. However, when anxiety-like and exploratory behavior were assessed via the ZM test (Fig 4E), no differences were seen in total distance travelled or percent time spent in the open arm of the apparatus (Fig 4F, G).

**Figure 4.**
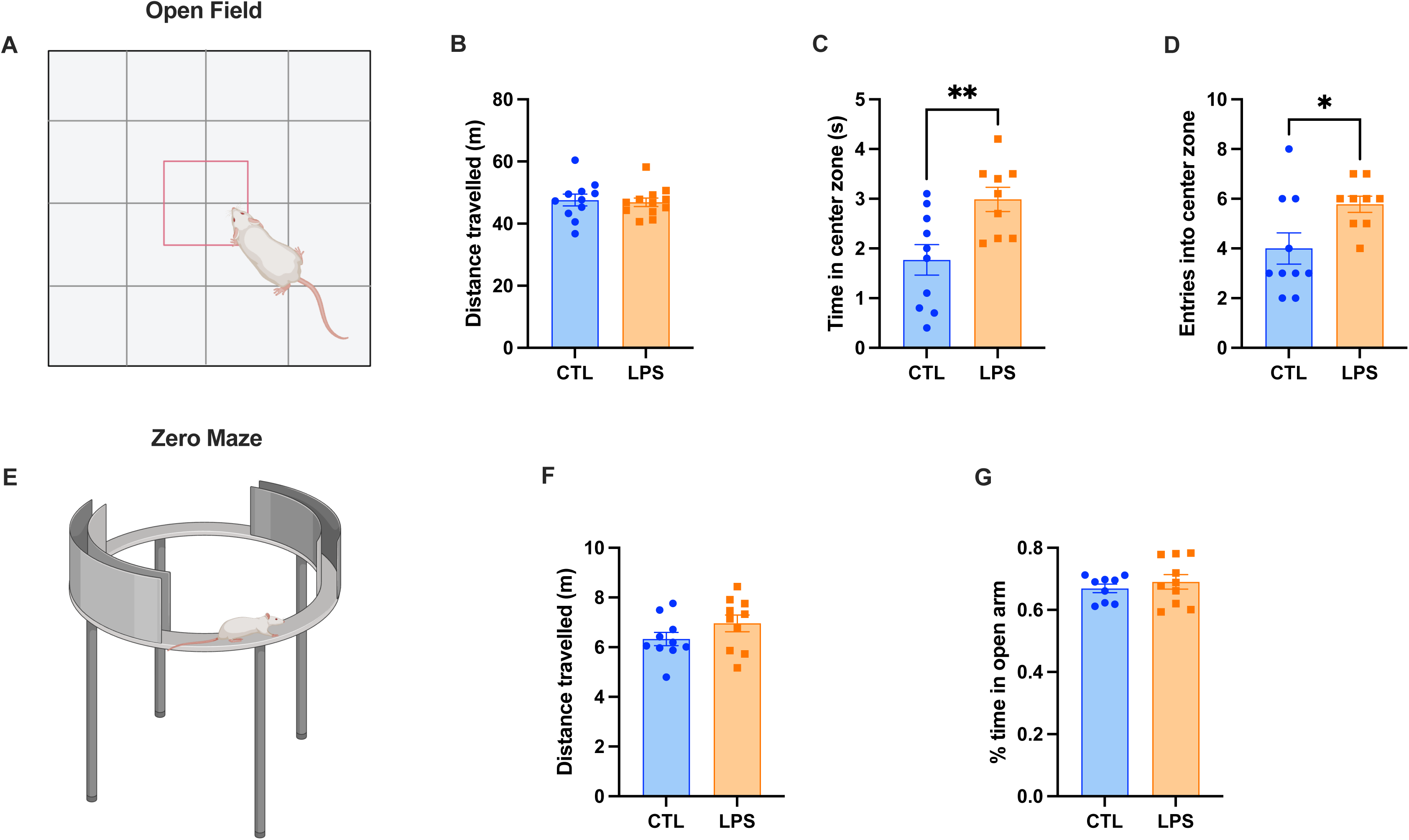
Chronic LPS administration decreases anxiety-like behavior. Anxiety-like behavior was significantly decreased in the open field test **(A)** with chronic LPS administration when compared to controls, with experimental rats spending more time **(C)** and making more entries into the center zone (**D**) when compared to control rats. Total distance travelled was unaffected **(B).** In the zero maze test (**E**), both total distance travelled **(F)** and percent time spent in the open arm of the apparatus **(G)** were unaffected. (LPS n = 9, CTL n = 10 for open field, LPS n = 10, CTL = 10 for zero maze; all between-subjects design; data are means ± SEM; *p<0.05, **p<0.01).

To assess whether impulsive responding is affected by LPS-mediated chronic inflammation, animals were tested in the DRL test (Fig 5A) to measure impulsive action. While there was no difference in overall learning during training (Fig 5B), LPS-treated rats made significantly fewer presses on the active lever, while earning the same number of rewards when compared to control rats on test day (Fig 5C, D). This led to LPS-treated rats having significantly higher efficiency in the task when compared to control animals (as measured by rewards earned over total active lever presses), indicating decreased impulsive responding (p < 0.05; Fig 5E).

**Figure 5.**
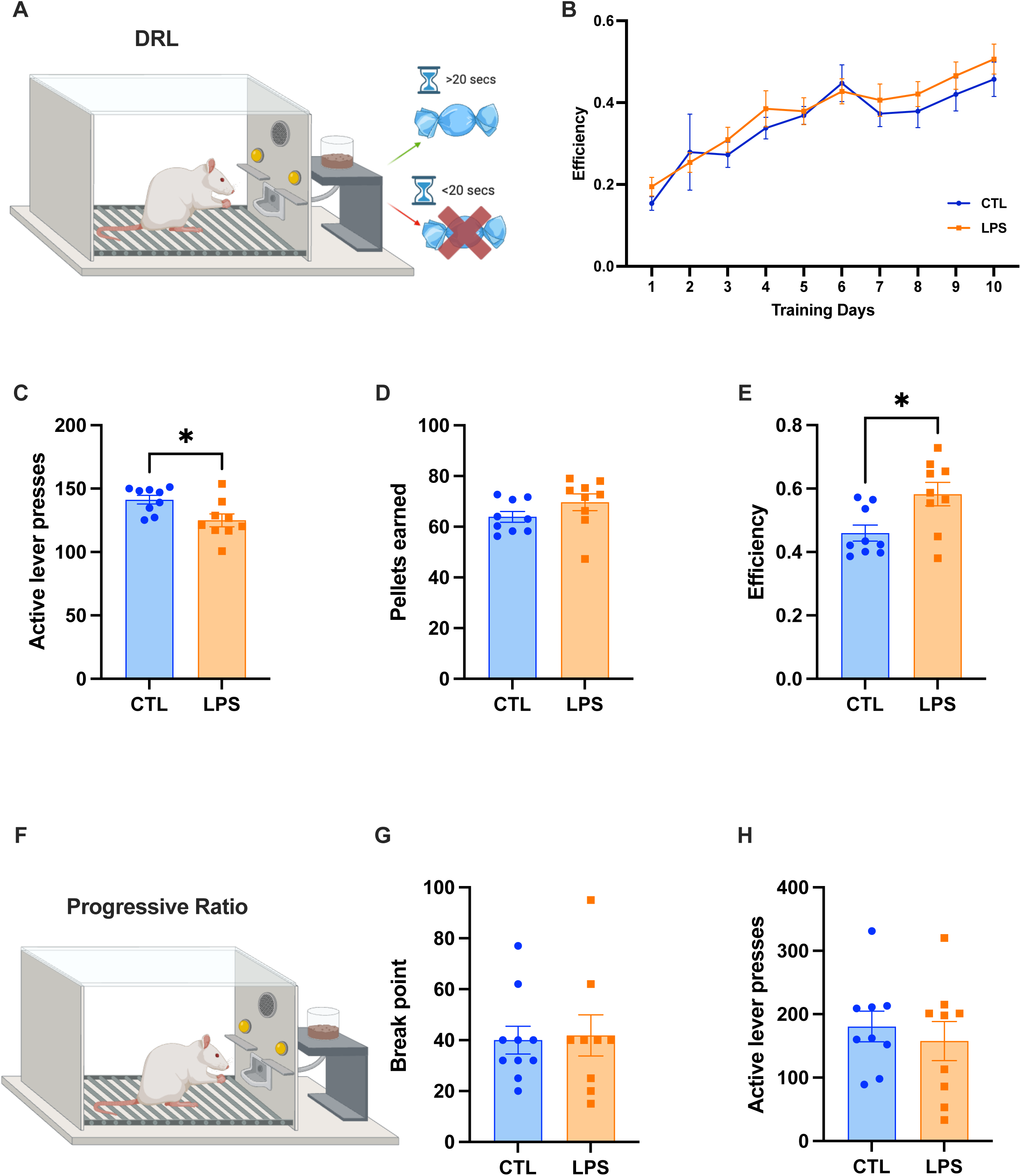
Chronic LPS administration decreases impulsive responding for palatable foods independently of reward motivation. In the DRL test of impulsive action **(A)**, animals were trained to wait at least 20s between lever presses in order to obtain a reward over the course of 10 days **(B)**. Results reveal that animals receiving chronic LPS administration made significantly fewer responses on the active lever **(C)**, while earning the same number of rewards **(D)**, leading to an increase in efficiency in the task **(E)** when compared to controls. Reward motivation as measured by the progressive ratio (PR) task **(F)** was unaffected, with both break point **(G)** and total active lever presses **(H)** remaining unaffected. (LPS n = 9 CTL = 9-10; all between-subjects design; data are means ± SEM; *p<0.05).

However, given that a decrease in responses on the active lever could indicate an overall difference in motivation to work for the reinforcement, animals were also tested in the progressive ratio (PR) operant task in which each successive reinforcer earned requires more lever presses than the previous reinforcer (Fig 5F), which is a common test of motivation to work for palatable food. There were no group differences in active lever presses as assessed in PR, nor were there any differences in the final breakpoint between LPS-treated and control rats (Fig 5G, H), suggesting that the effects on impulsive responding are not secondary to alterations in reward motivation.

## DISCUSSION

Obesity, Western diet consumption, and the concomitant increased gut permeability can lead to metabolic endotoxemia driving both chronic peripheral and central inflammation (6–8, 12). These factors independently are each associated with neurocognitive deficits, affecting cognitive domains such as anxiety, reward motivation, and learning and memory (15, 19–21, 23, 24). Thus, disentangling the causal relationships between these variables is increasingly difficult. Here, we modeled the chronic low-grade inflammation associated with obesity and/or Western diet consumption, yet in nonobese animals maintained on a healthy diet. Metabolic endotoxemia and its associated inflammatory profile was modeled using an indwelling mini osmotic pump to continuously dispense LPS solution. Our results reveal that low-dose LPS treatment over the course of 6 weeks is sufficient to significantly increase microglial recruitment in the amygdala (Amyg) and hippocampus (HPC; in both dorsal and ventral subregions), reduce serum corticosterone and melatonin, and decrease anxiety-like and impulsive behaviors. LPS treatment did not, however, affect astrocytic reactivity, HPC-dependent memory, or food reward-motivated operant responding.

Previous research indicates that obese individuals have higher levels of circulating LPS relative to healthy weight individuals (44), although the precise levels of serum LPS can differ greatly between studies (44). Some studies have considered LPS levels of 0-0.1 ng/mL to be physiologically relevant in mice (45), and serum LPS levels in our study fell within this range. One previous study has shown that individuals with obesity have roughly 1.6x higher LPS levels relative to their healthy weight counterparts (46), which is in line with the difference between saline- and LPS-treated animals in the present study. Thus, the present model appears to approximate physiological levels of LPS in circulation in obesity. Consistent with a previous study using the same dose of LPS in adult rats (29), LPS treatment in the present study produced an initial hypophagia and body weight loss, followed by a period of hyperphagia and weight regain until body weights were comparable to controls. In this previous study and with present results, the body weights of LPS-treated rats never exceeded that of controls, thus indicating that excessive body weight is not a characteristic in this model in rats. However, a previous study using adult mice and the same dose of chronic LPS revealed that experimental mice weighed significantly more than controls after 4 weeks of administration (7). Thus, while there may be species differences in the effects of chronic low-dose LPS with regards to body weight effects, results from the present study clearly model elevated LPS in circulation in the absence of obesity, and within a physiological range with regards to obesity.

Changes in behavioral responses may be driven by LPS-induced neuroinflammation (28, 47, 48). Chronic LPS exposure significantly increased microglial activation in the Amyg, dHPC, and vHPC, while astrocytic reactivity remained unaffected. Previous literature is consistent with these findings (48–50) and suggests that 6 weeks of LPS treatment is not sufficient time to develop significant alterations in astrocytic activity, but rather, up to 16 weeks of administration may be required to induce astrocytic activity (51). Microglial activation caused by LPS has been previously demonstrated in the HPC (47, 48), albeit with doses significantly higher than our own. Similarly, Huang et al. found increases in the size of microglia cell bodies in the basolateral amygdala following 3 acute injections of a supraphysiological dose of LPS (1mg/kg) 20 weeks prior to Iba1 analyses (52). Present results expand these previous findings to suggest that a low physiological dose of LPS when given chronically is sufficient to induce changes in microglia in the HPC and Amyg. These changes in microglia in key brains regions associated with several cognitive domains (i.e. learning, memory, exploratory behavior, food motivation) lead us to investigate possible behavioral outcomes associated with neuroinflammation.

While previous studies suggest that hippocampal-dependent memory is strongly impacted by neuroinflammation (27, 53–55), learning and memory were unaffected by LPS exposure in our study in the MWM, NLR, and NOIC procedures. The lack of differences in learning and memory was surprising as previous research has shown that LPS exposure can have deleterious effects on memory (47, 51, 56, 57). These discrepancies could be explained by our study’s relatively low dose of LPS (12.5ug/kg/hr) compared to previous studies, as well as by differences in the mode and timing of LPS administration. For example, a majority of previous studies utilized acute injections of a high dose of LPS to induce inflammation, with doses ranging from 167-750ug/mL LPS (47, 56, 57). Only one of these studies found deficits in MWM in wildtype animals, using an acute, supraphysiological dose of 500ug/mL LPS (47). Others found impairments in MWM only in transgenic Alzheimer’s disease model mice (56), or in NOIC following an acute injection of a high dose of LPS (57). Schirmbeck and colleagues, who found impairments in NLR in LPS treated rats, did utilize a chronic administration design, although injections took place weekly for 16 weeks, and at a significantly higher dose (500ug/kg) than our own (51). Collectively, our results expand previous literature to suggest that, when chronic endotoxemia is within a physiological range, the learning and memory deficits commonly associated with obesity and Western diet consumption are most likely not driven by peripheral or central inflammation.

Given that the amygdala is heavily implicated in fear and anxiety (58), and that we found significant differences in amygdalar neuroinflammation and exploratory behavior in the NOIC procedure, we investigated the effects of LPS-driven inflammation on established tests of exploratory and anxiety-like behavior. While acute, high dose (500-830ug/mL) LPS exposure reliably induced anxiety-like behavior in previous studies, (28, 59) the effects of chronic LPS exposure at a physiologically-relevant dose are unclear. When measured in the OFT, LPS-treated animals in the present study spent significantly more time in the center of the arena and made more entries into the center zone when compared to their control counterparts, which is consistent with a previous report (54). This could be interpreted as decreased anxiety-like behavior and increased exploratory behavior, which is consistent with our data from the NOIC procedure revealing more time investigating objects overall in LPS-vs. saline-treated rats. However, when anxiety-like behavior was assessed via the ZM task, no differences were found in time spent in the open arm of the apparatus, an outcome consistent with previous findings (51, 60). There are several possibilities as to why behavior in our study was affected in the OFT but not ZM task. As metabolic endotoxemia can take up to 4 weeks to fully develop (7), it is important to note that our behavioral testing took place around or shortly after this time point, but before the pumps dispense all their solution by 6wks. Given this short experimental window for behavioral testing, behavior in the ZM was assessed before the OFT with ZM testing on day 25 post-surgery and OF testing 34 days after LPS treatment initiation. Thus, it is possible that the effects of the LPS-induced chronic inflammation were more robust when behavior was assessed in the OFT vs. the ZM test, which took place shortly before the 4-week mark from pump implantation. It is also the case that the OFT and ZM procedures differ in various parameters, including elevation from the floor, procedure temporal length, and test lighting conditions. Nevertheless, our collective results suggest that exploratory behavior is increased and anxiety-like behavior is reduced in animals exposed to LPS-mediated chronic low-grade inflammation within a physiological range.

Increased entries into the center zone in OFT are often interpreted as decreased anxiety, but this behavior may also be driven by increased impulsivity. Therefore, we assessed the effects of chronic LPS exposure on impulsive responding for palatable foods. Results revealed that impulsivity for palatable food-reinforced responding was also altered by LPS-treatment, with experimental animals being significantly more efficient in earning rewards in the DRL test of impulsive action when compared to controls (interpreted as less impulsivity). While LPS-exposed rats did make significantly fewer presses on the active lever in the DRL task, we found no differences in the number of active lever presses or the breakpoint threshold when assessed in the operant PR task. This suggests that our findings in DRL are unlikely to be based solely on overall difference in food reward motivation, and that chronic low-grade inflammation driven by LPS exposure is sufficient to decrease impulsivity. Previous research suggests that LPS exposure can have a dampening effect on reward-motivated behavior (61–64) and increase impulsive responding (65–67), though differences in dose, mode, and timing of LPS administration could all play a part. For example, Lasselin and colleagues found differences in reward behavior in the Effort Expenditure for Rewards Task in humans following a single LPS injection, with LPS driving healthy individuals to button press less frequently for high-effort tasks (62). However, this effect was related to sleepiness, with LPS increasing sleepiness significantly 3hrs post-injection, and no differences in reward sensitivity were found (62). Similarly, male wildtype mice made significantly fewer nose pokes in a low-effort, low reward vs. high-effort, high reward choice task following a single injection of a supraphysiological dose of LPS (33mg/kg) 24hrs prior to testing (61). Thus, these previous findings taken together with present results suggest that while acute high doses of LPS can reduce reward-motivated responses, the effects of chronic physiological LPS treatment have minimal impact on motivated behavior, and can even reduce reward-driven impulsivity.

Regarding impulsivity, previous findings showing that LPS increased impulsive responding may also be explained by acute vs. chronic administration, as well as other contributing factors. In several studies, acute injections were given early in life (65, 66), during developmental time periods that are known to be particularly sensitive to environmental insults (68, 69). While Russell and colleagues found elevated premature responding in the 5-choice serial reaction time test in LPS-treated animals, acute injections took place immediately prior to behavioral testing and at a dose intended to cause sickness behavior (150ug/mL) (67). Thus, it is unsurprising that there are behavioral differences between their study using a supraphysiologic dose and the present findings. Collectively, our results expand previous literature by revealing that while acute and high dose LPS treatment may reliably reduce reward-motivated responses and increase impulsivity, chronic endotoxemia-induced inflammation within a physiological range can have the opposite effect.

It is possible that our behavioral results may be explained, in part, by differences in stress responsivity between LPS- and saline-treated animals, as serum corticosterone and melatonin were both significantly decreased in LPS-treated experimental animals when compared to controls. While previous research has shown that LPS increases serum corticosterone levels, it is likely due to the acute injection of a high dose of LPS (1.5mg/kg) prior to serum collection (70). The hypothalamic-pituitary-adrenal (HPA) axis, which plays a crucial role in mediating responses to stress, is known to exhibit bidirectional effects on cortisol levels (71, 72), and previous research suggests that stressors can exert negative feedback effects on HPA axis activity by suppressing corticotropin releasing hormone (CRH) neuron activity (73). Additionally, cortisol responses to LPS injections are dampened in individuals with obesity (74). These results are in line with our current study, suggesting that altered HPA axis activity may be implicated in behavioral changes associated with chronic inflammation (71, 72, 75).

Melatonin, which was reduced in circulation in LPS-treated rats in the present study, also plays an important role in HPA axis regulation, and is known to inhibit neuroinflammation (76). Papp et al. found that acute injections of melatonin enhanced open arm exploration in the elevated plus maze task (77), suggesting a possible role for melatonin in reducing anxiety-like behavior. Importantly, previous research also suggests that melatonin acts to attenuate the degradation of tight-junction proteins caused by LPS treatment (78), and that anxiety induced by an acute injection of a high dose of LPS (250ug/kg) 3hrs before testing is attenuated with concomitant melatonin injections in rats (79). These results show a potentially important role for melatonin in the modulation of LPS-mediated inflammation induced cognitive changes. However, the seemingly complex interaction between LPS and melatonin signaling, as well as how acute vs. chronic LPS may differentially impact melatonin levels and anxiety-like behavior, requires additional mechanistic follow-up studies.

Here, our collective results support a model in which elevated serum LPS caused by chronic exposure to a physiologically-relevant dose of LPS, and the concomitant peripheral and central inflammation, leads to a dampened HPA axis response and hyposensitivity to stress. This HPA axis dysregulation then results in decreased serum corticosterone and melatonin, as well as reduced impulsive responding and anxiety-like behavior. Importantly, these endotoxemia-mediated physiological and behavioral changes in the absence of obesity or Western diet consumption do not appear to mediate the widely established effects of both obesity and Western diet on the dysregulation of hippocampal-dependent memory. Future research should investigate the effects of chronic low-dose LPS exposure on HPA axis reactivity to identify precise mechanisms linking chronic inflammation to behavioral outcomes, as well to better understand the effects of acute vs. chronic inflammation on neurocognition when inflammatory treatments are kept within a physiological range. Further, it is important to follow up on what, if not neuroinflammation, mediates Western diet- and obesity-associated learning and memory impairments.

## ACKNOWLEDGEMENTS

The authors would like to thank the Kanoski Lab undergraduate research assistants for their support with the behavioral experiments. Figure subpanels 1A, 3A, 3E, 3H, 4A, 4E, 5A, and 5F were created with assistance from Biorender.com. This work was supported by the following funding sources: National Institute of Diabetes and Digestive and Kidney Diseases grant DK123423 (SEK), Postdoctoral Ruth L. Kirschstein National Research Service Award from the National Institute on Aging F32AG077932 (AH), Ruth L. Kirschstein Predoctoral Individual National Research Service Award F31DK138777 (MK).

## References

1. CDC. About Obesity [Online]. Obesity: 2024. https://www.cdc.gov/obesity/php/about/index.html [15 Apr. 2025].

2. Moreira APB, Texeira TFS, Ferreira AB, Peluzio M do CG, Alfenas R de CG. Influence of a high-fat diet on gut microbiota, intestinal permeability and metabolic endotoxaemia. British Journal of Nutrition 108: 801–809, 2012. doi: 10.1017/S0007114512001213.

3. Mishra SP, Wang B, Jain S, Ding J, Rejeski J, Furdui CM, Kitzman DW, Taraphder S, Brechot C, Kumar A, Yadav H. A mechanism by which gut microbiota elevates permeability and inflammation in obese/diabetic mice and human gut.

4. Rainone V, Schneider L, Saulle I, Ricci C, Biasin M, Al-Daghri NM, Giani E, Zuccotti GV, Clerici M, Trabattoni D. Upregulation of inflammasome activity and increased gut permeability are associated with obesity in children and adolescents. Int J Obes 40: 1026–1033, 2016. doi: 10.1038/ijo.2016.26.

5. Nagpal R, Newman TM, Wang S, Jain S, Lovato JF, Yadav H. Obesity-Linked Gut Microbiome Dysbiosis Associated with Derangements in Gut Permeability and Intestinal Cellular Homeostasis Independent of Diet. Journal of Diabetes Research 2018: 3462092, 2018. doi: 10.1155/2018/3462092.

6. Fuke N, Nagata N, Suganuma H, Ota T. Regulation of Gut Microbiota and Metabolic Endotoxemia with Dietary Factors. Nutrients 11: 2277, 2019. doi: 10.3390/nu11102277.

7. Cani PD, Amar J, Iglesias MA, Poggi M, Knauf C, Bastelica D, Neyrinck AM, Fava F, Tuohy KM, Chabo C, Waget A, Delmée E, Cousin B, Sulpice T, Chamontin B, Ferrières J, Tanti J-F, Gibson GR, Casteilla L, Delzenne NM, Alessi MC, Burcelin R. Metabolic Endotoxemia Initiates Obesity and Insulin Resistance. Diabetes 56: 1761– 1772, 2007. doi: 10.2337/db06-1491.

8. Pendyala S, Walker JM, Holt PR. A high-fat diet is associated with endotoxemia that originates from the gut. Gastroenterology 142: 1100–1101.e2, 2012. doi: 10.1053/j.gastro.2012.01.034.

9. Cani PD, Bibiloni R, Knauf C, Waget A, Neyrinck AM, Delzenne NM, Burcelin R. Changes in Gut Microbiota Control Metabolic Endotoxemia-Induced Inflammation in High-Fat Diet–Induced Obesity and Diabetes in Mice. Diabetes 57: 1470–1481, 2008. doi: 10.2337/db07-1403.

10. Peng X, Luo Z, He S, Zhang L, Li Y. Blood-Brain Barrier Disruption by Lipopolysaccharide and Sepsis-Associated Encephalopathy. Front Cell Infect Microbiol 11: 768108, 2021. doi: 10.3389/fcimb.2021.768108.

11. Brown GC. The endotoxin hypothesis of neurodegeneration. Journal of Neuroinflammation 16: 180, 2019. doi: 10.1186/s12974-019-1564-7.

12. Guillemot-Legris O, Muccioli GG. Obesity-Induced Neuroinflammation: Beyond the Hypothalamus. Trends in Neurosciences 40: 237–253, 2017. doi: 10.1016/j.tins.2017.02.005.

13. Anand SS, Friedrich MG, Lee DS, Awadalla P, Després JP, Desai D, de Souza RJ, Dummer T, Parraga G, Larose E, Lear SA, Teo KK, Poirier P, Schulze KM, Szczesniak D, Tardif J-C, Vena J, Zatonska K, Yusuf S, Smith EE, Canadian Alliance of Healthy Hearts and Minds (CAHHM) and the Prospective Urban and Rural Epidemiological (PURE) Study Investigators. Evaluation of Adiposity and Cognitive Function in Adults. JAMA Network Open 5: e2146324, 2022. doi: 10.1001/jamanetworkopen.2021.46324.

14. Smith E, Hay P, Campbell L, Trollor JN. A review of the association between obesity and cognitive function across the lifespan: implications for novel approaches to prevention and treatment. Obesity Reviews 12: 740–755, 2011. doi: 10.1111/j.1467-789X.2011.00920.x.

15. Kanoski SE, Davidson TL. Western Diet Consumption and Cognitive Impairment: Links to Hippocampal Dysfunction and Obesity. Physiol Behav 103: 59–68, 2011. doi: 10.1016/j.physbeh.2010.12.003.

16. Noble EE, Hsu TM, Kanoski SE. Gut to Brain Dysbiosis: Mechanisms Linking Western Diet Consumption, the Microbiome, and Cognitive Impairment. Front Behav Neurosci 11: 9, 2017. doi: 10.3389/fnbeh.2017.00009.

17. Noble EE, Olson CA, Davis E, Tsan L, Chen Y-W, Schade R, Liu C, Suarez A, Jones RB, de La Serre C, Yang X, Hsiao EY, Kanoski SE. Gut microbial taxa elevated by dietary sugar disrupt memory function. Transl Psychiatry 11: 194, 2021. doi: 10.1038/s41398-021-01309-7.

18. Hayes AMR, Lauer LT, Kao AE, Sun S, Klug ME, Tsan L, Rea JJ, Subramanian KS, Gu C, Tanios N, Ahuja A, Donohue KN, Décarie-Spain L, Fodor AA, Kanoski SE. Western diet consumption impairs memory function via dysregulated hippocampus acetylcholine signaling. Brain Behav Immun 118: 408–422, 2024. doi: 10.1016/j.bbi.2024.03.015.

19. Francis H, Stevenson R. The longer-term impacts of Western diet on human cognition and the brain. Appetite 63: 119–128, 2013. doi: 10.1016/j.appet.2012.12.018.

20. Cordner ZA, Tamashiro KLK. Effects of high-fat diet exposure on learning & memory. Physiology & Behavior 152: 363–371, 2015. doi: 10.1016/j.physbeh.2015.06.008.

21. Fulton S, Décarie-Spain L, Fioramonti X, Guiard B, Nakajima S. The menace of obesity to depression and anxiety prevalence. Trends in Endocrinology & Metabolism 33: 18–35, 2022. doi: 10.1016/j.tem.2021.10.005.

22. Volkow ND, Wang G-J, Baler RD. Reward, dopamine and the control of food intake: implications for obesity. Trends in Cognitive Sciences 15: 37–46, 2011. doi: 10.1016/j.tics.2010.11.001.

23. Wallace CW, Fordahl SC. Obesity and dietary fat influence dopamine neurotransmission: exploring the convergence of metabolic state, physiological stress, and inflammation on dopaminergic control of food intake. Nutr Res Rev 35: 236–251, 2022. doi: 10.1017/S0954422421000196.

24. Shi H, Schweren LJS, Ter Horst R, Bloemendaal M, van Rooij D, Vasquez AA, Hartman CA, Buitelaar JK. Low-grade inflammation as mediator between diet and behavioral disinhibition: A UK Biobank study. Brain Behav Immun 106: 100–110, 2022. doi: 10.1016/j.bbi.2022.07.165.

25. Lavagnino L, Arnone D, Cao B, Soares JC, Selvaraj S. Inhibitory control in obesity and binge eating disorder: A systematic review and meta-analysis of neurocognitive and neuroimaging studies. Neuroscience & Biobehavioral Reviews 68: 714–726, 2016. doi: 10.1016/j.neubiorev.2016.06.041.

26. Giel K, Teufel M, Junne F, Zipfel S, Schag K. Food-Related Impulsivity in Obesity and Binge Eating Disorder—A Systematic Update of the Evidence. Nutrients 9: 1170, 2017. doi: 10.3390/nu9111170.

27. de Paula GC, Brunetta HS, Engel DF, Gaspar JM, Velloso LA, Engblom D, de Oliveira J, de Bem AF. Hippocampal Function Is Impaired by a Short-Term High-Fat Diet in Mice: Increased Blood–Brain Barrier Permeability and Neuroinflammation as Triggering Events. Front Neurosci 15: 734158, 2021. doi: 10.3389/fnins.2021.734158.

28. Zheng Z-H, Tu J-L, Li X-H, Hua Q, Liu W-Z, Liu Y, Pan B-X, Hu P, Zhang W-H. Neuroinflammation induces anxiety- and depressive-like behavior by modulating neuronal plasticity in the basolateral amygdala. Brain Behav Immun 91: 505–518, 2021. doi: 10.1016/j.bbi.2020.11.007.

29. de La Serre CB, de Lartigue G, Raybould HE. Chronic exposure to Low dose bacterial lipopolysaccharide inhibits leptin signaling in vagal afferent neurons. Physiology & Behavior 139: 188–194, 2015. doi: 10.1016/j.physbeh.2014.10.032.

30. Broadbent NJ, Squire LR, Clark RE. Spatial memory, recognition memory, and the hippocampus. Proc Natl Acad Sci U S A 101: 14515–14520, 2004. doi: 10.1073/pnas.0406344101.

31. Clark RE, Broadbent NJ, Squire LR. Hippocampus and Remote Spatial Memory in Rats. Hippocampus 15: 260–272, 2005. doi: 10.1002/hipo.20056.

32. Balderas I, Rodriguez-Ortiz CJ, Salgado-Tonda P, Chavez-Hurtado J, McGaugh JL, Bermudez-Rattoni F. The consolidation of object and context recognition memory involve different regions of the temporal lobe. Learn Mem 15: 618–624, 2008. doi: 10.1101/lm.1028008.

33. Martínez MC, Villar ME, Ballarini F, Viola H. Retroactive interference of object-in-context long-term memory: Role of dorsal hippocampus and medial prefrontal cortex. Hippocampus 24: 1482–1492, 2014. doi: 10.1002/hipo.22328.

34. Suarez AN, Hsu TM, Liu CM, Noble EE, Cortella AM, Nakamoto EM, Hahn JD, de Lartigue G, Kanoski SE. Gut vagal sensory signaling regulates hippocampus function through multi-order pathways. Nat Commun 9: 2181, 2018. doi: 10.1038/s41467-018-04639-1.

35. Tsan L, Chometton S, Hayes AMR, Klug ME, Zuo Y, Sun S, Bridi L, Lan R, Fodor AA, Noble EE, Yang X, Kanoski SE, Schier LA. Early life low-calorie sweetener consumption disrupts glucose regulation, sugar-motivated behavior, and memory function in rats.

36. Hayes AMR, Tsan L, Kao AE, Schwartz GM, Décarie-Spain L, Tierno Lauer L, Klug ME, Schier LA, Kanoski SE. Early Life Low-Calorie Sweetener Consumption Impacts Energy Balance during Adulthood. Nutrients 14: 4709, 2022. doi: 10.3390/nu14224709.

37. Davis EA, Wald HS, Suarez AN, Zubcevic J, Liu CM, Cortella AM, Kamitakahara AK, Polson JW, Arnold M, Grill HJ, de Lartigue G, Kanoski SE. Ghrelin Signaling Affects Feeding Behavior, Metabolism, and Memory through the Vagus Nerve. Current Biology 30: 4510–4518.e6, 2020. doi: 10.1016/j.cub.2020.08.069.

38. Noble EE, Wang Z, Liu CM, Davis EA, Suarez AN, Stein LM, Tsan L, Terrill SJ, Hsu TM, Jung A-H, Raycraft LM, Hahn JD, Darvas M, Cortella AM, Schier LA, Johnson AW, Hayes MR, Holschneider DP, Kanoski SE. Hypothalamus-hippocampus circuitry regulates impulsivity via melanin-concentrating hormone. Nat Commun 10: 4923, 2019. doi: 10.1038/s41467-019-12895-y.

39. Décarie-Spain L, Liu CM, Lauer LT, Subramanian K, Bashaw AG, Klug ME, Gianatiempo IH, Suarez AN, Noble EE, Donohue KN, Cortella AM, Hahn JD, Davis EA, Kanoski SE. Ventral hippocampus-lateral septum circuitry promotes foraging-related memory. Cell Reports 40, 2022. doi: 10.1016/j.celrep.2022.111402.

40. Hsu TM, Hahn JD, Konanur VR, Lam A, Kanoski SE. Hippocampal GLP-1 Receptors Influence Food Intake, Meal Size, and Effort-Based Responding for Food through Volume Transmission. Neuropsychopharmacol 40: 327–337, 2015. doi: 10.1038/npp.2014.175.

41. Bankhead P, Loughrey MB, Fernández JA, Dombrowski Y, McArt DG, Dunne PD, McQuaid S, Gray RT, Murray LJ, Coleman HG, James JA, Salto-Tellez M, Hamilton PW. QuPath: Open source software for digital pathology image analysis. Sci Rep 7: 16878, 2017. doi: 10.1038/s41598-017-17204-5.

42. Schneider CA, Rasband WS, Eliceiri KW. NIH Image to ImageJ: 25 years of image analysis. Nat Methods 9: 671–675, 2012. doi: 10.1038/nmeth.2089.

43. Jimenez JC, Su K, Goldberg AR, Luna VM, Biane JS, Ordek G, Zhou P, Ong SK, Wright MA, Zweifel L, Paninski L, Hen R, Kheirbek MA. Anxiety Cells in a Hippocampal-Hypothalamic Circuit. Neuron 97: 670–683.e6, 2018. doi: 10.1016/j.neuron.2018.01.016.

44. Gnauck A, Lentle RG, Kruger MC. Chasing a ghost? – Issues with the determination of circulating levels of endotoxin in human blood. Critical Reviews in Clinical Laboratory Sciences 53: 197–215, 2016. doi: 10.3109/10408363.2015.1123215.

45. Guo S, Al-Sadi R, Said HM, Ma TY. Lipopolysaccharide Causes an Increase in Intestinal Tight Junction Permeability in Vitro and in Vivo by Inducing Enterocyte Membrane Expression and Localization of TLR-4 and CD14. The American Journal of Pathology 182: 375–387, 2013. doi: 10.1016/j.ajpath.2012.10.014.

46. Trøseid M, Nestvold TK, Rudi K, Thoresen H, Nielsen EW, Lappegård KT. Plasma Lipopolysaccharide Is Closely Associated With Glycemic Control and Abdominal Obesity: Evidence from bariatric surgery. Diabetes Care 36: 3627–3632, 2013. doi: 10.2337/dc13-0451.

47. Zhao J, Bi W, Xiao S, Lan X, Cheng X, Zhang J, Lu D, Wei W, Wang Y, Li H, Fu Y, Zhu L. Neuroinflammation induced by lipopolysaccharide causes cognitive impairment in mice. Sci Rep 9: 5790, 2019. doi: 10.1038/s41598-019-42286-8.

48. Zhang J, Zhang N, Lei J, Jing B, Li M, Tian H, Xue B, Li X. Fluoxetine shows neuroprotective effects against LPS-induced neuroinflammation via the Notch signaling pathway. International Immunopharmacology 113: 109417, 2022. doi: 10.1016/j.intimp.2022.109417.

49. Llorens-Martín M, Jurado-Arjona J, Fuster-Matanzo A, Hernández F, Rábano A, Ávila J. Peripherally triggered and GSK-3β-driven brain inflammation differentially skew adult hippocampal neurogenesis, behavioral pattern separation and microglial activation in response to ibuprofen. Transl Psychiatry 4: e463–e463, 2014. doi: 10.1038/tp.2014.92.

50. Hill JD, Zuluaga-Ramirez V, Gajghate S, Winfield M, Sriram U, Persidsky Y. Activation of GPR55 induces neuroprotection of hippocampal neurogenesis and immune responses of neural stem cells following chronic, systemic inflammation. Brain Behav Immun 76: 165–181, 2019. doi: 10.1016/j.bbi.2018.11.017.

51. Schirmbeck GH, Seady M, Fróes FT, Taday J, Da Ré C, Souza JM, Gonçalves CA, Leite MC. Long-term LPS systemic administration leads to memory impairment and disturbance in astrocytic homeostasis. Neurotoxicology 99: 322–331, 2023. doi: 10.1016/j.neuro.2023.11.009.

52. Huang H-T, Chen P-S, Kuo Y-M, Tzeng S-F. Intermittent peripheral exposure to lipopolysaccharide induces exploratory behavior in mice and regulates brain glial activity in obese mice. Journal of Neuroinflammation 17: 163, 2020. doi: 10.1186/s12974-020-01837-x.

53. Czerniawski J, Guzowski JF. Acute Neuroinflammation Impairs Context Discrimination Memory and Disrupts Pattern Separation Processes in Hippocampus. J Neurosci 34: 12470–12480, 2014. doi: 10.1523/JNEUROSCI.0542-14.2014.

54. Ryan SM, Nolan YM. Neuroinflammation negatively affects adult hippocampal neurogenesis and cognition: can exercise compensate? Neuroscience & Biobehavioral Reviews 61: 121–131, 2016. doi: 10.1016/j.neubiorev.2015.12.004.

55. Barrientos RM, Kitt MM, Watkins LR, Maier SF. Neuroinflammation in the normal aging hippocampus. Neuroscience 309: 84–99, 2015. doi: 10.1016/j.neuroscience.2015.03.007.

56. Valero J, Mastrella G, Neiva I, Sánchez S, Malva JO. Long-term effects of an acute and systemic administration of LPS on adult neurogenesis and spatial memory. Front Neurosci 8: 83, 2014. doi: 10.3389/fnins.2014.00083.

57. Czerniawski J, Miyashita T, Lewandowski G, Guzowski JF. Systemic lipopolysaccharide administration impairs retrieval of context–object discrimination, but not spatial, memory: Evidence for selective disruption of specific hippocampus-dependent memory functions during acute neuroinflammation. *Brain*, Behavior, and Immunity 44: 159–166, 2015. doi: 10.1016/j.bbi.2014.09.014.

58. Davis M. The Role of the Amygdala in Fear and Anxiety.

59. Bassi GS, Kanashiro A, Santin FM, de Souza GEP, Nobre MJ, Coimbra NC. Lipopolysaccharide-Induced Sickness Behaviour Evaluated in Different Models of Anxiety and Innate Fear in Rats. Basic & Clinical Pharmacology & Toxicology 110: 359– 369, 2012. doi: 10.1111/j.1742-7843.2011.00824.x.

60. Loh MK, Stickling C, Schrank S, Hanshaw M, Ritger AC, Dilosa N, Finlay J, Ferrara NC, Rosenkranz JA. Liposaccharide-induced sustained mild inflammation fragments social behavior and alters basolateral amygdala activity. Psychopharmacology 240: 647–671, 2023. doi: 10.1007/s00213-023-06308-8.

61. Vichaya EG, Hunt SC, Dantzer R. Lipopolysaccharide Reduces Incentive Motivation While Boosting Preference for High Reward in Mice. Neuropsychopharmacol 39: 2884–2890, 2014. doi: 10.1038/npp.2014.141.

62. Lasselin J, Treadway MT, Lacourt TE, Soop A, Olsson MJ, Karshikoc B, Paues-Göranson S, Axelsson J, Dantzer R, Lekander M. Lipopolysaccharide Alters Motivated Behavior in a Monetary Reward Task: a Randomized Trial. Neuropsychopharmacol 42: 801–810, 2017. doi: 10.1038/npp.2016.191.

63. De La Garza R, Asnis GM, Fabrizio KR, Pedrosa E. Acute diclofenac treatment attenuates lipopolysaccharide-induced alterations to basic reward behavior and HPA axis activation in rats. Psychopharmacology 179: 356–365, 2005. doi: 10.1007/s00213-004-2053-x.

64. Huwart SJP, Fayt C, Gangarossa G, Luquet S, Cani PD, Everard A. TLR4-dependent neuroinflammation mediates LPS-driven food-reward alterations during high-fat exposure. J Neuroinflammation 21: 305, 2024. doi: 10.1186/s12974-024-03297-z.

65. Gruzdeva VA, Sharkova AV, Zaichenko MI, Grigoryan GA. Effects of Early Proinflammatory Stress on Manifestations of Impulsive Behavior in Rats of Different Ages and Sexes. Neurosci Behav Physi 51: 1079–1085, 2021. doi: 10.1007/s11055-021-01168-1.

66. Straley ME, Van Oecelen W, Theze S, Sullivan AM, O’Mahony SM, Cryan JF, O’Keece GW. Distinct alterations in motor & reward seeking behavior are dependent on the gestational age of exposure to LPS-induced maternal immune activation. *Brain*, Behavior, and Immunity 63: 21–34, 2017. doi: 10.1016/j.bbi.2016.06.002.

67. Russell B, Hrelja KM, Adams WK, Zeeb FD, Taves MD, Kaur S, Soma KK, Winstanley CA. Differential effects of lipopolysaccharide on cognition, corticosterone and cytokines in socially-housed vs isolated male rats. Behavioural Brain Research 433: 114000, 2022. doi: 10.1016/j.bbr.2022.114000.

68. Hocman DJ, Powell TL, Barrett ES, Hardy DB. Developmental origins of metabolic diseases. Physiological Reviews 101: 739–795, 2021. doi: 10.1152/physrev.00002.2020.

69. Van den Bergh BRH, van den Heuvel MI, Lahti M, Braeken M, de Rooij SR, Entringer S, Hoyer D, Roseboom T, Räikkönen K, King S, Schwab M. Prenatal developmental origins of behavior and mental health: The influence of maternal stress in pregnancy. Neuroscience & Biobehavioral Reviews 117: 26–64, 2020. doi: 10.1016/j.neubiorev.2017.07.003.

70. Girard-Joyal O, Faragher A, Bradley K, Kane L, Hrycyk L, Ismail N. Age and sex differences in c-Fos expression and serum corticosterone concentration following LPS treatment. Neuroscience 305: 293–301, 2015. doi: 10.1016/j.neuroscience.2015.06.035.

71. Rutters F, La Fleur S, Lemmens S, Born J, Martens M, Adam T. The Hypothalamic-Pituitary-Adrenal Axis, Obesity, and Chronic Stress Exposure: Foods and HPA Axis. Curr Obes Rep 1: 199–207, 2012. doi: 10.1007/s13679-012-0024-9.

72. Rusch JA, Layden BT, Dugas LR. Signalling cognition: the gut microbiota and hypothalamic-pituitary-adrenal axis. Front Endocrinol 14, 2023. doi: 10.3389/fendo.2023.1130689.

73. Jiang Z, Chen C, Weiss GL, Fu X, Stelly CE, Sweeten BLW, Tirrell PS, Pursell I, Stevens CR, Fisher MO, Begley JC, Harrison LM, Tasker JG. Stress-induced glucocorticoid desensitizes adrenoreceptors to gate the neuroendocrine response to somatic stress in male mice. Cell Rep 41: 111509, 2022. doi: 10.1016/j.celrep.2022.111509.

74. Lasselin J, Benson S, Hebebrand J, Boy K, Weskamp V, Handke A, Hasenberg T, Remy M, Föcker M, Unteroberdörster M, Brinkhoc A, Engler H, Schedlowski M. Immunological and behavioral responses to *in vivo* lipopolysaccharide administration in young and healthy obese and normal-weight humans. *Brain*, Behavior, and Immunity 88: 283–293, 2020. doi: 10.1016/j.bbi.2020.05.071.

75. Packard AEB, Egan AE, Ulrich-Lai YM. HPA axis-Interaction with Behavioral Systems. Compr Physiol 6: 1897–1934, 2016. doi: 10.1002/cphy.c150042.

76. Wang Y-Q, Jiang Y-J, Zou M-S, Liu J, Zhao H-Q, Wang Y-H. Antidepressant actions of melatonin and melatonin receptor agonist: Focus on pathophysiology and treatment. Behav Brain Res 420: 113724, 2022. doi: 10.1016/j.bbr.2021.113724.

77. Papp M, Litwa E, Gruca P, Mocaër E. Anxiolytic-like activity of agomelatine and melatonin in three animal models of anxiety. Behavioural Pharmacology 17: 9, 2006. doi: 10.1097/01.fbp.0000181601.72535.9d.

78. Hu Y, Wang Z, Pan S, Zhang H, Fang M, Jiang H, Zhang H, Gao Z, Xu K, Li Z, Xiao J, Lin Z. Melatonin protects against blood-brain barrier damage by inhibiting the TLR4/ NF-κB signaling pathway after LPS treatment in neonatal rats. Oncotarget 8: 31638– 31654, 2017. doi: 10.18632/oncotarget.15780.

79. Nava F, Carta G. Melatonin reduces anxiety induced by lipopolysaccharide in the rat. Neuroscience Letters 307: 57–60, 2001. doi: 10.1016/S0304-3940(01)01930-9.

